# The *Vibrio fischeri* type VI secretion system incurs a fitness cost under host-like conditions

**DOI:** 10.1101/2023.03.07.529561

**Authors:** Alecia N. Septer, Garrett Sharpe, Erika A. Shook

## Abstract

The type VI secretion system (T6SS) is an interbacterial weapon composed of thousands of protein subunits and predicted to require significant cellular energy to deploy, yet a fitness cost from T6SS use is rarely observed. Here, we identify host-like conditions where the T6SS incurs a fitness cost using the beneficial symbiont, *Vibrio fischeri*, which uses its T6SS to eliminate competitors in the natural squid host. We hypothesized that a fitness cost for the T6SS could be dependent on the cellular energetic state and used theoretical ATP cost estimates to predict when a T6SS-dependent fitness cost may be apparent. Theoretical energetic cost estimates predicted a minor relative cost for T6SS use in fast-growing populations (0.4-0.45% of total ATP used cell^-1^), and a higher relative cost (3.1-13.6%) for stationary phase cells. Consistent with these predictions, we observed no significant T6SS-dependent fitness cost for fast-growing populations typically used for competition assays. However, the stationary phase cell density was significantly lower in the wild-type strain, compared to a regulator mutant that does not express the T6SS, and this T6SS-dependent fitness cost was between 11 and 23%. Such a fitness cost could influence the prevalence and biogeography of T6SSs in animal-associated bacteria. While the T6SS may be required in kill or be killed scenarios, once the competitor is eliminated there is no longer selective pressure to maintain the weapon. Our findings indicate an evolved genotype lacking the T6SS would have a growth advantage over its parent, resulting in the eventual dominance of the unarmed population.

## Introduction

The type VI secretion system (T6SS) is a molecular syringe through which cells transport diverse effectors into neighboring cells or the extracellular environment (1). T6SSs are found broadly in Gram-negative bacterial genomes where they function in interbacterial killing and host interactions, thereby conferring a competitive advantage in polymicrobial systems (1). However, the fitness cost for T6SS-using populations remains understudied. Cells can harbor multiple T6SS structures per cell, each composed of ~4600 subunits (Fig 1A) (2, 3, 4). Moreover, several protein subunits are secreted with each “firing” and a ClpV ATPase is required to disassemble the contracted sheath, adding to the cost of each secretion event (2). Because cells have a limited amount of ATP to hydrolyze, and this ATP pool must be divided into biomass production (including T6SSs) and cellular maintenance (including T6SS firing) (Fig 1B), it has been predicted that T6SS use may be energetically costly (2, 5, 6, 7, 8, 9, 10, 11). Yet the cost of the T6SS has remained largely theoretical, with only one reported observation of a conditional fitness cost in *Campylobacter jejuni* (12). Thus, there is a need to identify when T6SSs incur a fitness cost in additional model organisms.

**Figure 1.**
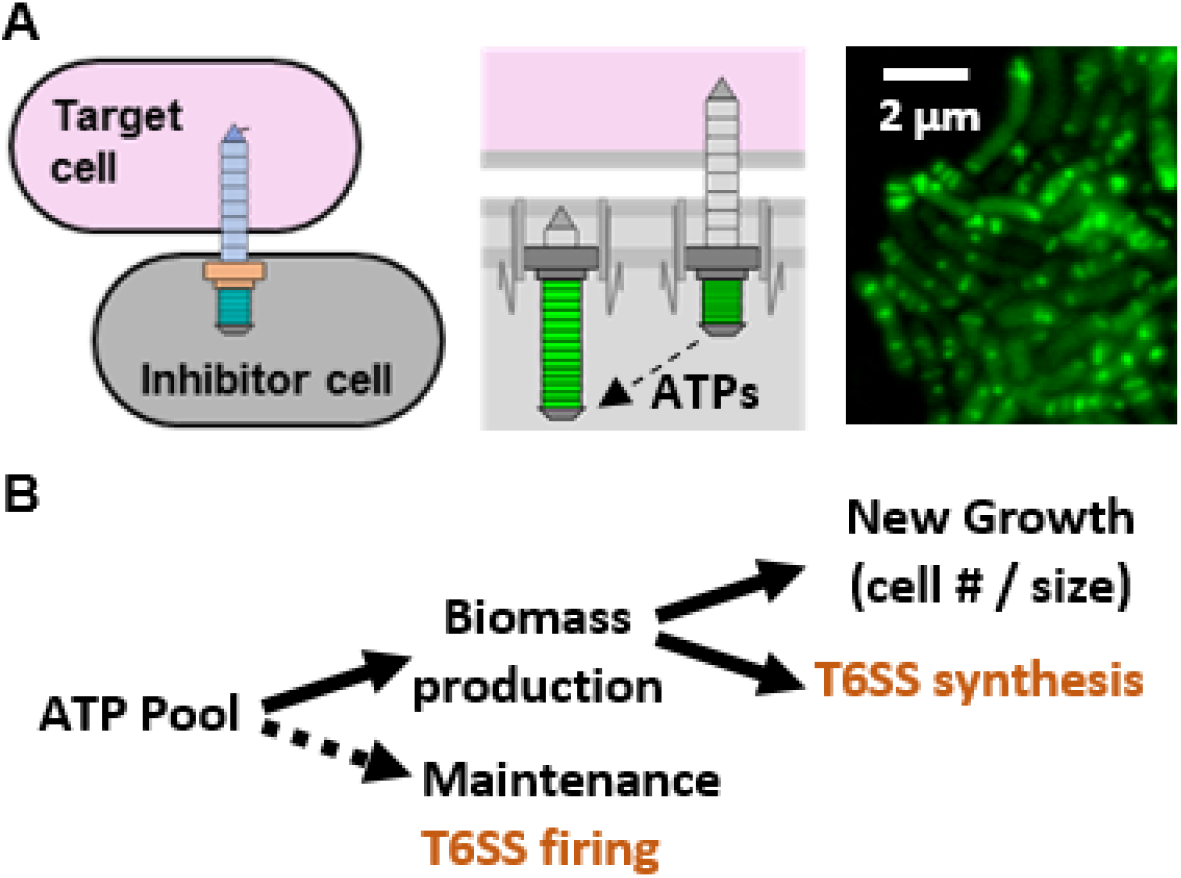
The T6SS is predicted to be a costly structure in stationary phase cells. (A) Diagram of T6SS structures in an inhibitor cell in both an extended (before firing] and contracted state (after firing]. The ClpV ATPase is required to disassemble the contracted sheath in order to build an extended sheath for firing. Representative fluorescence microscopy image of *V. fischeri* ES401 with a *vipA-gfp* expression vector to visualize T6SS sheaths in live cells. (B) Diagram of how ATP consumption feeds into biomass and maintenance of cell. Dashed line indicates primary energy use in stationary phase cells when population growth has arrested.

The beneficial marine symbiont, *Vibrio fischeri*, is a model organism for studying quorum sensing, host-colonization, and interbacterial competition (13). Although *V. fischeri* is the only species cultured from the light organs of wild-caught adult *Euprymna scolopes* squid, multiple strains are routinely isolated from an individual animal, indicating that these light organ communities are not clonal and contain multiple genotypes (14, 15). Previous work has shown that *V. fischeri* isolates have a strain-specific type VI secretion system on chromosome II (T6SS2) that is required for killing competitor strains in culture (16). The T6SS2 is conditionally expressed and requires a combination of host-like conditions and transcription factors encoded on the T6SS2-containing genomic island (GI) to activate expression and assembly of the weapon (17, 18, 19, 20, 21). Although T6SS2 is not required for *V. fischeri* to colonize the squid during clonal infection assays in the absence of a competitor, it is required to prevent cocolonized crypts in the juvenile light organ during competitive colonization assays (16, 22). Thus, the T6SS2 provides a competitive advantage during host colonization, yet the energetic cost of this advantage remains unknown.

Here, we combined theoretical energy cost estimates of the T6SS with culture-based experiments to identify possible T6SS-dependent fitness costs. We hypothesized that the relative T6SS fitness cost is dependent on the amount of available ATP, and therefore could be predicted using theoretical energy cost estimates that incorporate the cellular energetic state. We used the squid symbiont, *V. fischeri*, as a model organism to calculate the relative energy costs of T6SS and predict when it might incur a fitness cost. We chose *V. fischeri* strain ES401 because 1) it encodes the T6SS2 genomic island (GI) that is used to prevent co-colonized crypts within the squid host (16, 22), 2) T6SS2 expression can be controlled by varying culture conditions (18), and 3) we have a regulator mutant that is unable to activate T6SS2 expression (20). Thus, we can control when T6SS2 is expressed using both culture conditions and mutant strains. Using this approach we discovered that cells grown in T6SS2-activating conditions that mimic the host environment incur a fitness cost observable as reduced cell density in stationary phase. These findings, which reveal an ecologically-relevant fitness cost for the T6SS, expands the utility of *V. fischeri* as a model organism relevant to studying how a costly interbacterial weapon can be maintained in a natural system.

## Results

### Theoretical T6SS energy cost estimates

We calculated theoretical energy cost estimates for T6SS2 as the percent of ATP used per cell in fast-growing or stationary phase populations using the parameters outlined in the methods and Table 1. Briefly, calculations included the energy required to build all protein subunits at a cost of 4-5 ATP equivalents per peptide bond (23), as well as ATP-dependent ClpV cost estimates (2). We assumed each cell contains six T6SSs per cell (11), and each daughter cell must synthesize three new structures during active growth, while stationary cells only incur the cost of firing the T6SS (Fig 2). To calculate the percent of total ATP use, we divided the ATP cost estimates for T6SS2 cell^-1^ by the total ATPs used cell^-1^ over the same time scale using values reported in Deng *et al.*, (24), which calculated ATPs used cell^-1^ sec^-1^ for *E. coli* growth in exponential (6.4 million ATPs cell^-1^ sec^-1^) and stationary phase (200,000 - 800,000 ATPs cell^-1^ sec^-1^), and found that the cellular ATP consumption rate increased with increasing growth rate. Our calculations predict that the relative cost of T6SS2 in fast-growing cells is low (0.4 - 0.45%), but increases to 3.1 - 13.6% in stationary phase cells (Table 2), suggesting cells might incur a detectable fitness cost as cells transition to a lower cellular energy state.

**Figure 2.**
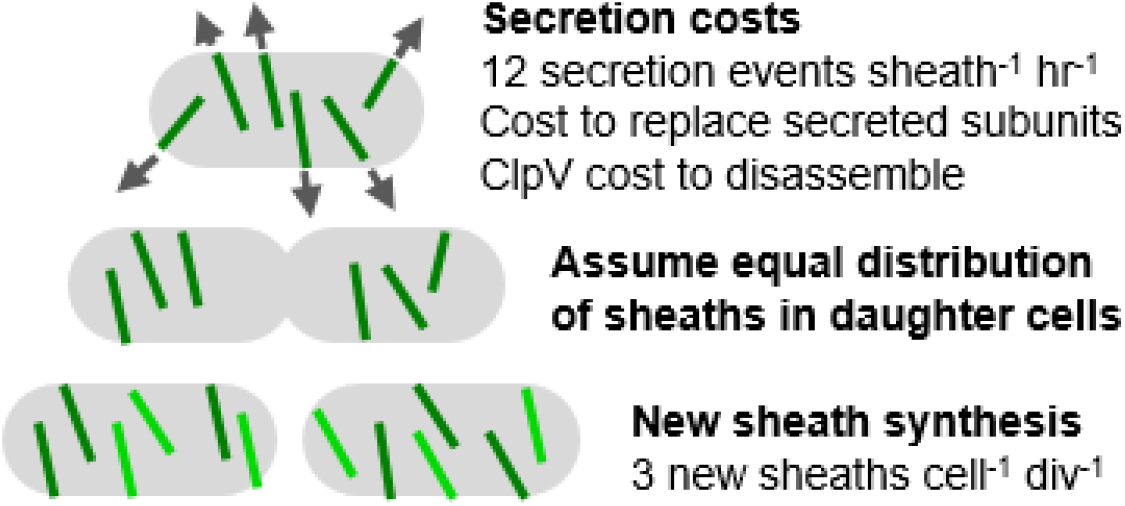
Parameters far ATP cast estimates far T6SS synthesis and secretion. ATP costs for T6SS secretion can be grouped into two categories: the cost to replace secreted protein subunits and ClpV ATPase activity to disassemble the contracted sheath (fast- and slow-growing cells, see shaded rows in Table 2),and the cost to synthesize new sheaths after cell division (fast-growing cells only. The total estimated ATP cost for T6SS per cell includes the sum of the cost to synthesize new sheaths, replace secreted subunits, and disassemble the contracted sheath over a defined time scale.

**Table 1.** Parameters for calculating estimated ATP cost of T6SS synthesis and use.

| Feature | Value | Empirical or Estimate | Reference(s) |
| --- | --- | --- | --- |
| Structures per cell | 6 | Empirical | Lin and Smith *et al.* 2021 |
| New sheaths made per division | 3 | Estimate (based on equal distribution of sheaths in daughter cells) | Lin and Smith *et al.* 2021 |
| Firings per hour per structure | 12 | Empirical based on other bacterial system | Basler *et al.* 2015 |
| Doubling time for *V. fischeri* (condition used here) | 4 hr | Empirical | This study |
| ATP equivalents for tRNA charging and polypeptide synthesis | 4-5 | Empirical | Calhoun and Swartz, 2007 |
| AAs per T6SS subunit | Varies | Empirical | Bultman *et al.*, 2019 |
| Number of subunits per structure | Varies | Empirical from other model bacterial species | Dix *et al.* 2018, Liebl *et al.* 2019, Basler 2015 |
| Number of subunits lost per T6SS firing | Varies | Estimates | Basler 2015 |
| ATPs used by ClpV per structure | 750,000 | Estimate | Basler 2015 |
| ATPs used per sec per cell | 200,000-800,000 (stationary);<br>6,400,000 (exponential) | Empirical (*E. coli*) | Deng *et al.* 2021 |

**Table 2.**
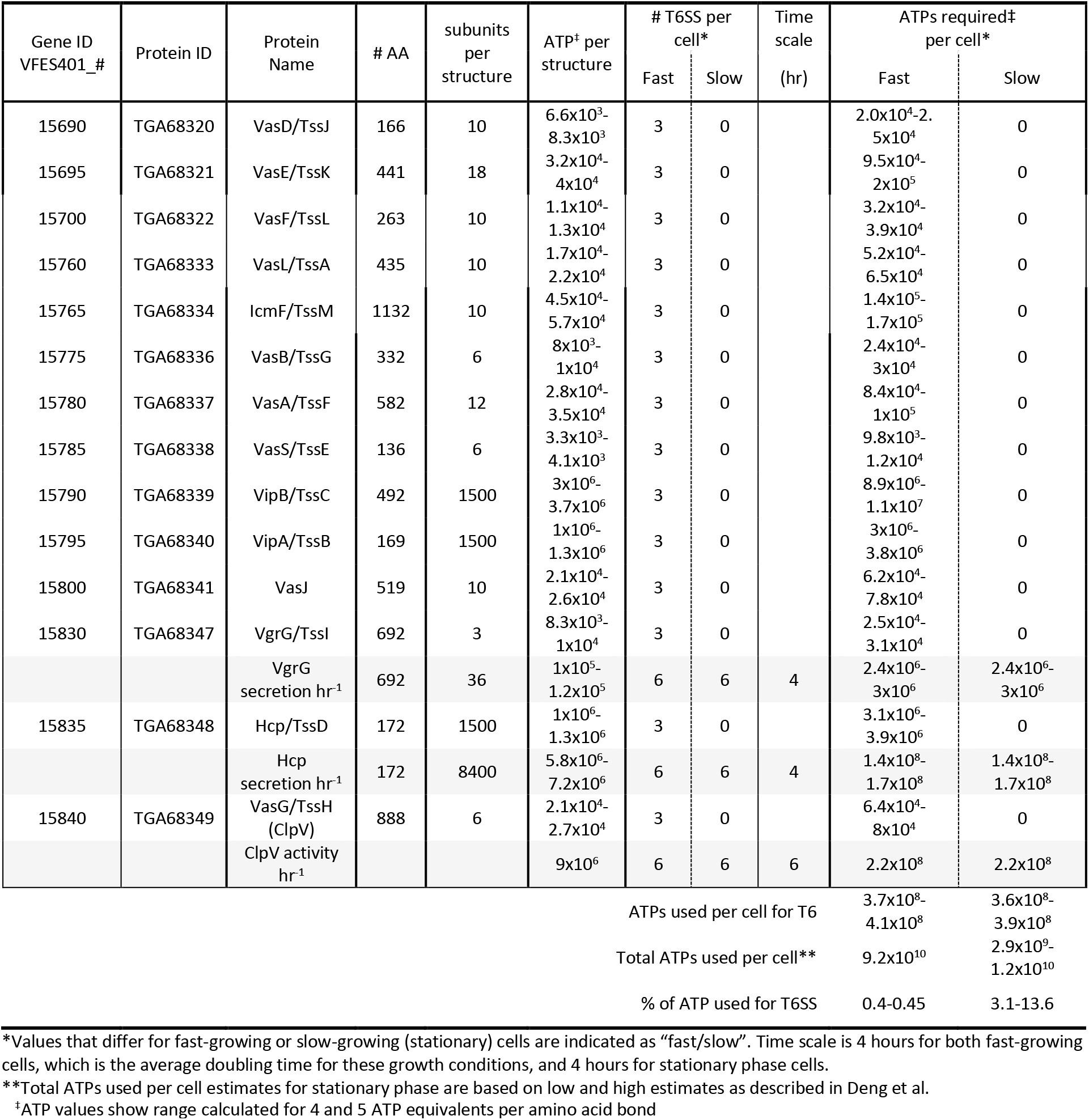
Theoretical ATP cost estimates for T6SS use in slow and fast-growing cells.

### Quantifying the T6SS-dependent fitness cost in host-like growth conditions

To test this prediction, we grew both wild-type *V. fischeri* strain ES401 and the *tasR* regulator mutant (strain SS51), which does not activate expression of the T6SS2 genes, in different media conditions to control T6SS2 expression. Previous work has shown that *V. fischeri* does not fully activate T6SS2 in liquid LBS (18); however, when cells are grown in a host-like high viscosity liquid hydrogel (LBS + PVP), T6SS2 expression is activated and abundant sheaths are observed in a *tasR*-dependent manner (20). When we grew the wild-type parent and *tasR* mutant in liquid LBS where T6SS2 is not induced, we did not see a difference in growth curves or growth rates in full-nutrient medium (LBS) or when we limited media nutrients by half (½ LBS) (Fig 3AB). When repeated in media supplemented with PVP to activate T6SS2 expression, we consistently observed a higher final optical density at stationary phase for the *tasR* mutant, compared to the wild type, in both LBS and ½ LBS hydrogel media (Fig 3C). There was a slight, but not significant, difference in growth rates between the parent and *tasR* mutant strains (Fig 3D), suggesting any apparent fitness cost in these fast-growing populations was too small to distinguish based on the rate of change in optical density. When we calculated the percent fitness cost by comparing the final optical densities (Fig 3E), there was no fitness cost in the absence of the T6SS2-inducing conditions (LBS) but we calculated a statistically significant fitness cost of 11-23% in the T6SS2-inducing conditions (PVP). These values are similar to the theoretical percent energy cost estimates for stationary phase cells (Table 2).

**Figure 3.**
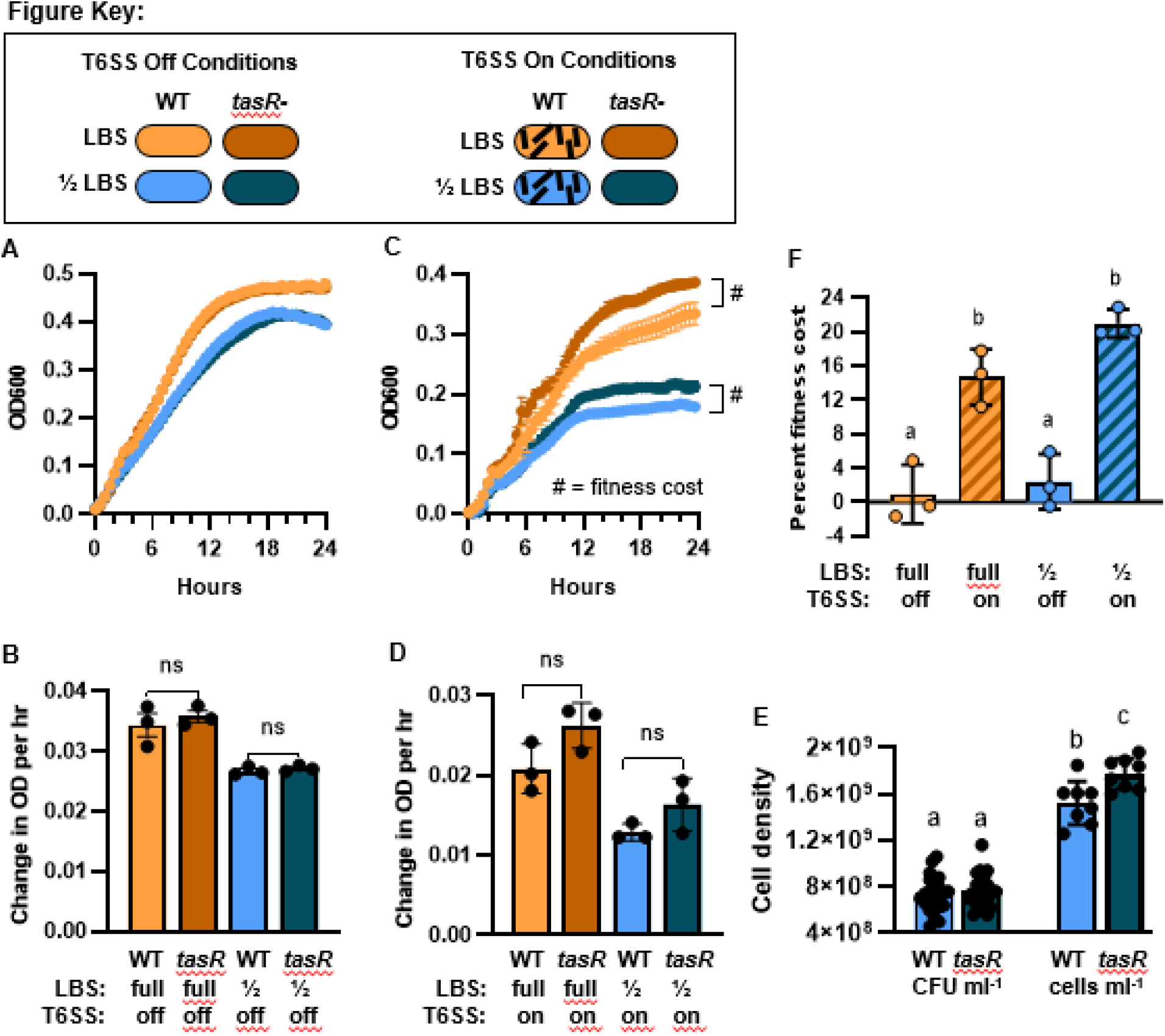
A significant T6S5-de pendent fitness cost is observed for stationary phase cell density. (A-D) Data from growth curves for wild-type *V. fischeri* ES401 [light shades) and the *tasR* mutant strain SS5l [dark shades) grown in a plate reader in LBS (orange shades) or ½ LBS liquid (brown shades) media without (A-B) or with (C-D) 5%PVP tc activate T6SS2 expression in wild-type cells. Data show OD6600 readings over time (A and C; error bars show standard error) and growth rates between hours 3 and 8 (B and D; error bars show standard deviation). Experiments were performed at east three times with biological triplicates, “ns” indicates not significant based on a t-test comparing growth rates for WT and *tasR* mutants within each growth condition. (E) The percent fitness costfor final OD from panels A and C. Letters indicate statistical significance (p<0.05) using a One-Way ANOVA and Bartlett’s corrected p-values. (F) CFU and direct cell counts in stationary phase cultures (16-24 hr) for WT and *tasR* mutant cultures grown in ½ LBS + PVP under the same conditions used for panel B. Letters indicate statistical significance (p<0.01) using a One-Way ANOVA and Tuke/s corrected p-values. Graph shows combined data from two independent experiments, each with at least four biological replicates. Error bars show standard deviation.

To confirm whether the higher optical density in the *tasR* mutant cultures was indeed due to higher cell numbers, we first quantified colony forming units (CFUs) for stationary-phase wild-type and *tasR* mutant cultures grown in ½ LBS with PVP. The average CFU ml^-1^ was 7.4 × 10^9^ and 7. 6 × 10^9^ for wildtype and *tasR* mutant, respectively, and was not significantly different (Fig 3F). Because CFU counts are indirect measurements of viable cell numbers that rely on cells to grow into a colony, and thus could underestimate the cells in a stationary phase sample, we quantified cell numbers using a flow cytometer. Direct cell counts confirmed that 1) viable cell count estimates using CFU values are lower than what is observed for direct cell counts that include both live and dead cells expected in stationary phase and, 2) the *tasR* mutant cultures contained on average 16.6% more cells than the wild-type cultures (Fig 3F).

## Discussion

In this work, we aimed to use energetic cost estimates for the T6SS to predict when a fitness cost might be most apparent for the model organism, *Vibrio fischeri*. Although this study identified a conditional T6SS-dependent fitness cost that correlates with predicted energy cost estimates, the underlying mechanisms of this cost remain unknown and will require further exploration. It could be that, without the ATP cost required to build and fire T6SSs, the *tasR* mutant cells may be able to divert additional energy into biomass, compared to the wild-type parent that has expended ATP on T6SS2, resulting in the slightly higher final cell density for *tasR* mutant cultures. The T6SS2-dependent fitness cost could be a result of the cumulative effect of the small, but not statistically significant, slower growth rate during fast growth (Fig 3D), and/or it could be due to the higher relative cost of the T6SS as cells transition into stationary phase. Moreover, other indirect costs of the T6SS that are independent of ATP use may impact population numbers, such as the recently-described cell death observed in clonal colonies of T6SS-producing *Vibrio cholerae* cells (25). Although this T6SS-mediated self-killing has not been observed for *V. fischeri* colonies on agar plates, it has yet to be explored in liquid hydrogel media. Despite these unknowns, this work suggests that selective pressures maintain T6SS2 in the majority of *V. fischeri* genomes (26) in spite of its fitness cost, underscoring its importance in competing for host colonization sites.

This work also revealed important details to consider when aiming to quantify a T6SS2-dependent fitness cost in *V. fischeri*. Past work has shown that there is no apparent fitness cost for a structural T6SS2 mutant, specifically *vasA_2/tssF_2*, for multiple strains grown in culture or during host colonization assays (16, 18, 22). However, it is worth noting that these experiments used either optical density readings during fast growth and/or CFU counts to determine cell numbers, approaches that we show here are not sufficient to detect the T6SS2-dependent fitness cost. Although CFU counts are excellent ways to determine viable cell counts during competition assays, which aim to determine how many viable cells remain in a population that can divide and expand that population’s genotype, it is an indirect count for overall cell numbers. Because the T6SS2-dependent fitness cost may not be detected with CFU counts, future work examining any potential fitness cost in the host, or in host-like culture conditions, may require quantification methods using optical density and/or direct cell counts.

It is also unknown whether *V. fischeri* maintain active T6SSs during stationary phase. Previous experiments visualizing sheaths in live cells examined cells from 12 hour cultures or earlier (16, 17, 18, 19, 20, 22), prior to transition into stationary phase. It is also unknown when or if the T6SS2 is turned off after it is activated, as earlier work in *V. fischeri* focused on identifying the inducing conditions and timescales for T6SS2 activation (17, 18, 19). Given that stationary phase is when rates of cell death and cell division are equal, it will be important to use approaches that quantify the T6SS2 on/off state at the single-cell level. Future work exploring the extent to which the T6SS2 weapon is maintained or decommissioned in stationary phase cells will greatly inform our understanding of the fitness cost and how such regulatory strategies may impact competitive outcomes.

Despite these unknowns, recent work that integrates subcellular T6SS dynamics into an agent-based model for *V. fischeri* probed the impact of the cost of this weapon in silico (11). Lin and Smith *et al*. competed strains that both wield T6SS weapons in an integrated agent-based model where the per cell sheath number and rate of sheath synthesis incurred a growth penalty. The simulation revealed a tipping point where cells that quickly produce many sheaths paid a high cost that prevented their own growth, resulting in a competitor with a smaller arsenal overtaking the heavily-armed population. These in silico findings, combined with our results, underscore the need for cells to strike a balance between the energetic cost of a T6SS and the fitness advantage it can confer. Although the T6SS is necessary to prevent a population from being eliminated by a competitor, too costly of a weapon can be self-defeating.

This study also provides a possible mechanism as to why some of the *V. fischeri* strains isolated from wild-caught adult squid are lacking a functional T6SS2 (16, 26), despite its clear advantage during initial colonization. Previous work suggested two possible explanations that are not necessarily mutually exclusive (16): 1) juvenile squid are initially colonized with genotypes lacking a functional T6SS2 that proliferate in crypts that are monocolonized or cocolonized by other T6SS2-deficient strains, and/or 2) initial crypt communities begin with multiple genotypes that require a functional T6SS2 to persist and exclude competitive genotypes, and over the lifetime of the squid the T6SS2 GI is lost or acquires a mutation deactivating T6SS2 expression because the selective pressure to maintain this weapon is removed once a co-colonizing competitor is eliminated. However, the latter hypothesis could only be plausible if there were a fitness cost associated with the T6SS2 in *V. fischeri*, such that a cell that acquires a T6SS2-off mutation would eventually outgrow the parent strain to become a dominate genotype in the light organ, which is later recovered by researchers.

This work reveals a fitness cost for T6SS2 and given the magnitude of the cost, it is conceivable that a T6SS2-off mutation occurring in symbiotic populations early during colonization of the animal would provide sufficient time for the T6SS2-off genotype to dominate and be selected during cultivation from adult animals. Indeed, if we assume a single mutant cell with a T6SS2-off mutation arises within a population of 10^6^ T6SS2-on parent cells, and the T6SS2-off genotype no longer incurs the 20 percent fitness cost per day, based on data in Fig 3, then the T6SS2-off genotype could theoretically become equally abundant as the T6SS2-on parent within two months, and become the dominant genotype by three months (Fig 4). If such growth dynamics occur in the squid, one might expect to isolate more T6SS2-on genotypes when sampling the light organs of juveniles, compared to adults. Indeed, recent work has shown the potential for *V. fischeri* to genetically diversify within the host light organ (27), and a metagenomics study in human guts described a significantly higher proportion of *Bacteroides fragilis* strains contained the GA3 T6SS genotype in infant gut microbiomes, compared to adults (28), suggesting the interplay between fitness costs and selective pressure may influence the T6SS prevalence in animal-associated bacteria across diverse hosts.

**Figure 4.**
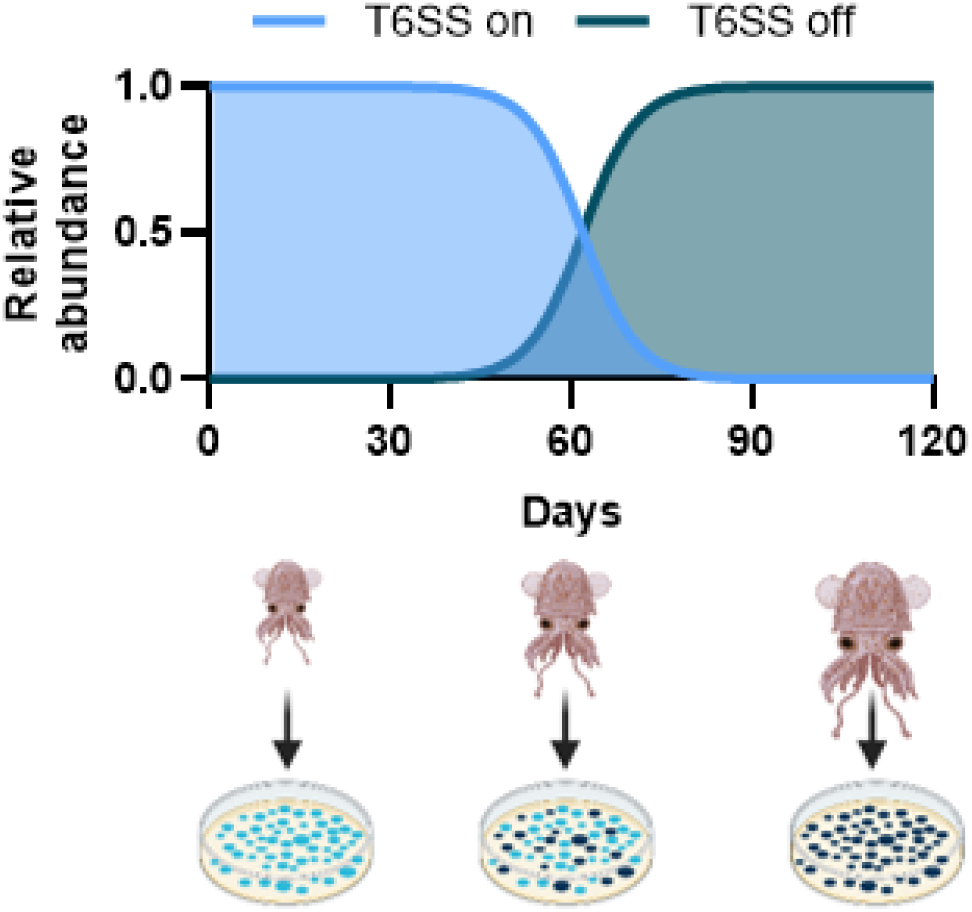
Model for how T6SS cost could alter the prevelance of T6SS genotypes in the host. Estimated relative abundance for T6SS-on genotype and a T6SS-off mutant that emerged at day 0 as a single cell in a population of 10^6^ T6SS-on parent cells. Change in relative abundance is based on the accumulation of 20 percent fitness cost per day reported in Fig 3F. Image made using BioRender.com.

In summary, this work revealed several important findings regarding the fitness cost associated with the T6SS. First, the relative cost is predicted to be higher in lower energy states. Therefore, it may be informative to investigate the potential fitness cost of T6SSs in other species when cells are in stationary phase or under growth conditions that may reduce the cellular ATP pool. Second, we have established a way to quantify the T6SS fitness cost in *V. fischeri*, which can be used to perform in vitro and in vivo evolution experiments to better understand how selective pressures maintain or decommission a costly weapon. Such studies are needed to interpret the genotypic variation observed for T6SS prevalence in natural microbiomes.

## Materials and Methods

### Strains and growth conditions

*Vibrio fischeri* strain ES401 is the wild-type strain used in this study, which was isolated from an adult *Euprymna scolopes* light organ (29). This work also used ES401 mutant strain SS51, which is a previously isolated transposon mutant with a disruption in the *tasR* regulator gene that has reduced expression of T6SS proteins (20). Bacteria were grown in Luria-Bertani with salts (LBS), or ½ LBS (half of LBS nutrients added) media without or with 5% (w/vol) of polyvinylpyrrolidone (PVP360, Sigma-Aldrich). To make LBS broth: 950 mL DI water, 50 mL 1M Tris-HCl pH 7.5, 20 g NaCl, 10 g tryptone, 5 g yeast extract. To make ½ LBS broth 5 g L^-1^ and 2.5 g L^-1^ were used for tryptone and yeast extract, respectively. To make PVP hydrogel media, broth media was first made and autoclaved and allowed to cool to 70°C, 5% w/vol PVP was added and allowed to dissolve in 70°C water bath. Hydrogel media was stored two weeks at room temperature and used within two weeks.

### Growth curves

To assess fitness cost in the form of growth dynamics wild-type ES401 or the *tasR* mutant (SS51) were grown at 200 rpm shaking and 24°C overnight in LBS medium and diluted 1:100 into 200 μl LBS or ½ LBS broth without or with 5% PVP in a clear 96-well plate. OD_600_ values were measured every 30 minutes using a Tecan plate reader over 24 hours at 24°C using a kinetic run, and the OD_600_ values for each treatment were blanked with an average OD_600_ value from either uninoculated ½ LBS broth with 5% PVP or ½ LBS without 5% PVP. Growth curves were performed at least three times with at least three biological replicates per strain and treatment.

### Flow cytometry enumeration cell counts

After 24 hours of the kinetic run experiment, 100 μL of each replicate for the ½ LBS with and without 5% PVP treatments for ES401 and SS51, as well as the ½ LBS plus and minus 5% PVP blanks, were diluted 5-fold in 400 μL of 36 psu NaCl solution and vortexed at max speed for 5 minutes to break up aggregates into single cells. The 5-fold dilutions of each replicate were then diluted 20-fold and then an additional 100-fold in 36 psu NaCl solution for a final dilution of 10,000x. For each 10,000x diluted replicate, 190 μL was mixed with 10 μL of 20X SYBER Green I nucleic acid stain (Lonza Walkersville Inc, Walkersville, MD) in a well of a transparent 96-well plate and incubated in the dark for 15 minutes before being placed in the Guava easyCyte HT flow cytometer (488 nm, 50 mW laser). The instrument settings were as follows: measurements of a sample continued until 2 minutes of measurement or 30,000 observations with the Grn-B threshold set to 25. The gating was set such that observations with fluorescence measurements between 2×10^2^ to 7×10^4^ and side scatter (SSC-HLog) between 15 and 3×10^3^ were counted as cells. Flow cytometry results were then blank and dilution corrected.

### Theoretical ATP energetic cost estimates

To calculate the estimated ATP cost of building and using the T6SS, we performed cost estimates for both fast- and slow-growing populations. For fast-growing cells, we assume six T6SS structures per cell (11), where cells must build three new T6SS structures after cell division and all six structures fire at a rate of 12 secretion events hr^-1^ structure^-1^ (2). For stationary phase cells where growth and death rates are balanced, we assume no net T6SS structure synthesis and only calculate the estimated cost of T6SS firing for six structures cell^-1^. First, we calculated the ATP cost for the synthesis of a new T6SS structure based on the number of each structural subunits, and a cost of 4-5 ATP equivalents for each amino acid added to a growing polypeptide chain, where 2 ATP-equivalents are required to charge the tRNA and 2-3 ATP equivalents are required to move the peptide chain through the ribosome for amino acid addition (23, 30, 31, 32). Calculations for protein synthesis cost estimates were performed for 4 and 5 ATP equivalents, resulting in a range of ATP cost estimates. We next added the estimated cost of synthesis of protein subunits that are lost and must be replaced during active secretion, including Hcp and VgrG components (2). Finally, we included ATP cost estimates for disassembly of the fired structure in order to rebuild a sheath for firing. The ClpV protein has been predicted to require as much as 750,000 ATPs per structure disassembly (2), although these values have not been experimentally validated and may represent a high estimate. For a complete list of calculation parameters, including references and whether the units are estimates or empirical, see Table 1.

Equation 1 is used to calculate the estimated ATP cost of T6SS cell^-1^ by summing the ATP cost for each protein subunit, where *n* is the number of protein subunits, in this case 14 (Table 2), and the number of events hr^-1^ refers to either the number of cell divisions (0.25 hr^-1^) or secretions (12 hr^-1^). For each subunit, the entire ATP cost estimate is then multiplied by the time scale of four hours (the time for the optical density to double during fast growth).

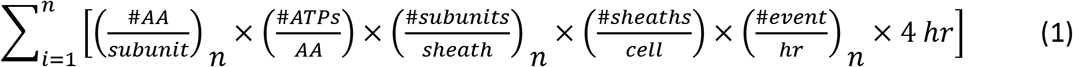

Equation 2 is used to calculate the estimated ATPs cell^-1^ used over the same timescale used in equation 1 (4 hours). The estimated total ATPs used cell^-1^ sec^-1^ is taken from Deng *et al*. (24), which reported 200,000-800,000 ATPs used cell^-1^ sec^-1^ for cells in stationary phase (i.e. “slow” in Table 2) and 6,400,000 ATPs used cell^-1^ sec^-1^ for cells in exponential phase (i.e. “fast” in Table 2).

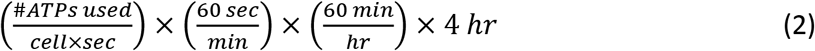

Equation 1 (ATPs used for T6SS cell^-1^) and equation 2 (ATPs used cell^-1^) can be used to estimate the percent of ATP used for T6SS as follows: 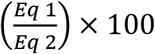

### Calculating fitness cost

To calculate the fitness cost from growth curves we used the following equation using the final OD_600_ values: [(SS51_OD600_ – WT_OD600_)/SS51_OD600_] x 100, where SS51 is the ES401 *tasR* mutant that does not activate expression of T6SS2 components and WT is the ES401 parent strain that activates T6SS expression in the presence of PVP.

## Acknowledgements

The authors thank Drs. Marc Alperin, Scott Gifford, and Marek Basler for helpful discussions, and Stephanie Smith for the microscopy image in Figure 1. ANS and EAS were supported on NIH NIGMS grant R35 GM137886; GS was supported by Gordon and Betty Moore Foundation Grant GBMF9328.

